# HIV-1 Gag specifically restricts PI(4,5)P2 and cholesterol mobility in living cells creating a nanodomain platform for virus assembly

**DOI:** 10.1101/556308

**Authors:** C. Favard, J. Chojnacki, P. Merida, N. Yandrapalli, J. Mak, C. Eggeling, D. Muriaux

## Abstract

HIV-1 Gag protein self-assembles at the plasma membrane of infected cells for viral particle formation. Gag targets lipids, mainly the phosphatidylinositol (4, 5) bisphosphate, at the inner leaflet of this membrane. Here, we address the question whether Gag is able to trap specifically PI(4,5)P2 or other lipids during HIV-1 assembly in the host CD4+ T lymphocytes. Lipid dynamics within and away from HIV-1 assembly sites was determined using super-resolution STED microscopy coupled with scanning Fluorescence Correlation Spectroscopy in living T cells. Analysis of HIV-1 infected cells revealed that, upon assembly, HIV-1 is able to specifically trap PI(4,5)P2, and cholesterol, but not phosphatidylethanolamine or sphingomyelin. Furthermore, our data show that Gag is the main driving force to restrict PI(4,5)P2 and cholesterol mobility at the cell plasma membrane. This is first direct evidence showing that HIV-1 creates its own specific lipid environment by selectively recruiting PI(4,5)P2 and cholesterol, as a membrane nano-platform for virus assembly.

## Introduction

During HIV-1 replication cycle, the assembly and release of newly made viral particles occurs at the plasma membrane of the infected cells (Fig 1a). Budded HIV-1 particles display an unusual lipid membrane composition that differs significantly from that of the plasma membrane of the host cell. It is enriched in sphingomyelins (SPMs), glycosphingolipids, cholesterol (Chol), and phosphoinositides, such as phosphatidylinositol (4, 5) bisphosphate lipid (PI(4,5)P2)(1, 2) and characterized by highly ordered, i.e. densely packed, lipid organization (1, 3). This raises the question on how virus components interact with lipids in the plasma membrane of the host cells to gather such specific lipid environment during assembly. Previous studies of HIV-lipid interactions were mainly based on in vitro model systems (4–8), tagged phospholipid interacting protein domains (9), or virus lipidomics (1, 2, 4). However, direct observations of the relevant molecular interactions in the living cells have for long been challenged by the limited spatial resolution of conventional optical microscopy. Fortunately, recent advances in super-resolution microscopy techniques have enabled for characterization of molecular mobility at the nanoscale, i.e. on the scale of individual (< 140 nm) HIV-1 particles. For example, experiments employing a combination of super-resolution STED microscopy and fluorescence correlation spectroscopy (STED-FCS (10) or scanning STED-FCS, sSTED-FCS (11, 12)) revealed a low mobility of molecules on HIV-1 surface (3) due to the high degree of lipid packing in the virus membrane (12, 13). The retroviral protein Gag is thought to be a key player in altering the HIV-1 lipid environment as it is not only the main structural determinant of the particle assembly, but it also mediates interactions between assembling virus particle and plasma membrane lipids. HIV-1 Gag is composed of 4 domains, MA, CA, NC and p6 and 2 spacer peptides (sp1 and sp2). The MA domain of Gag is myristoylated on its N-terminus and contains a highly basic region (named HBR), both responsible for Gag anchoring and targeting to the cell plasma membrane (PM)(14). The CA domain is involved in Gag-Gag interaction for multimerization. The NC domain is required for genomic RNA encapsidation and also for Gag-Gag multimerization through RNA (15). The p6, and part of the NC domain, recruit ESCRT cell factors for particle budding (16). The space peptide sp1 is also required for proper particle assembly and budding (17). The binding of Gag to the plasma membrane is primarily resulting from a bipartite signal within MA (reviewed in (14)). It is understood that MA is thought to interact with the acidic lipids of the PM inner leaflet(18), although recent studies suggest that the NC domain of Gag could also play a role(19). Substitutions of the MA basic residues have shown to affect targeting and membrane binding of HIV-1 Gag both in liposomes (20) and in cells (21). Furthermore, a lipidomic study has suggested that Gag interacts with acidic phospholipids in cells since virions produced by HIV-1 infected cells were enriched in PI(4,5)P2 via a MA dependent manner (2), and this interpretation is supported by recent report using tunable PI(4,5)P2 level to track Gag anchoring onto cell membrane via imaging analysis (22). The unusual lipid contents in HIV show that HIV particles require a specific lipid membrane environment for HIV-1 assembly. A key question in HIV assembly is whether Gag is targeted toward pre-existing lipid domains at the plasma membrane, or whether Gag actively creates its unique lipid compositions for virus formation. Historically, Gag has been shown to associate with the detergent resistant membrane (DRM) fractions (23, 24), which are enriched in SPM and Chol, and rafts properties of virion-cholesterol impact on virus function (25, 26). Saad et al.(27) have used truncated acyl chains PI(4,5)P2 to show the unsaturated sn2 acyl chain of the PI(4,5)P2 is trapped in an hydrophobic pocket while the myristate embed into the lipid membrane, leaving the complex MA:PI(4,5)P2 to partition to more liquid ordered membrane environments. Using phase separated giant unilamellar vesicles (GUV), Keller et al., have shown that multimerizable MA are partitioning into more liquid disordered membrane environments (6). In addition, using coarse grained molecular dynamics, researchers have highlighted that HIV-1 MA interacts only with PI(4,5)P2 sugar head and that PI(4,5)P2 is concentrated within the immediate vicinity of MA (28). Recent NMR studies on HIV-1 MA in interactions with lipidic membranes confirmed these results (29). Finally, our recent lipidic membrane-based study demonstrated that recombinant HIV-1 Gag was able to segregate PI(4,5)P2, and cholesterol, but excluded SPM upon Gag multimerization, and that Gag preferred to partition into PI(4,5)P2 enriched liquid disordered lipid membrane environments on GUV rather than liquid ordered lipid membrane environments (7). Together, these results suggested that HIV1 Gag may generate its own PI(4,5)P2 enriched membrane environment at the cell plasma membrane rather than assembling on pre-existing areas or domains with distinct lipid membrane environments. This is consistent with a study utilizing a novel biological tool to tune PI(4,5)P2 level at the cell plasma membrane, recently reported that PI(4,5)P2 not only targets Gag to the cell membrane but it is necessary for assembly site formation (22). In order to study this dynamic equilibrium of Gag-lipid interaction in living host CD4+ T cells, here we employ super-resolution sSTED-FCS approach to analyse lipid mobility inside and outside HIV-1 assembly sites on fluorescent lipid labelled CD4 T cells infected by HIV-1 or expressing Gag only. We find that infectious HIV1 is able to immobilize PI(4,5)P2, and Chol, but not phosphatidylethanolamine (PE) and SPM in these host cells, and that only Gag is required to restrict the movement of these lipid molecules at the assembly sites. Our results highlight that HIV-1 Gag selectively trap PI(4,5)P2 and Cholesterol in host CD4 T cells to create its own specialized lipid membrane environment for virus assembly.

**Fig. 1.**
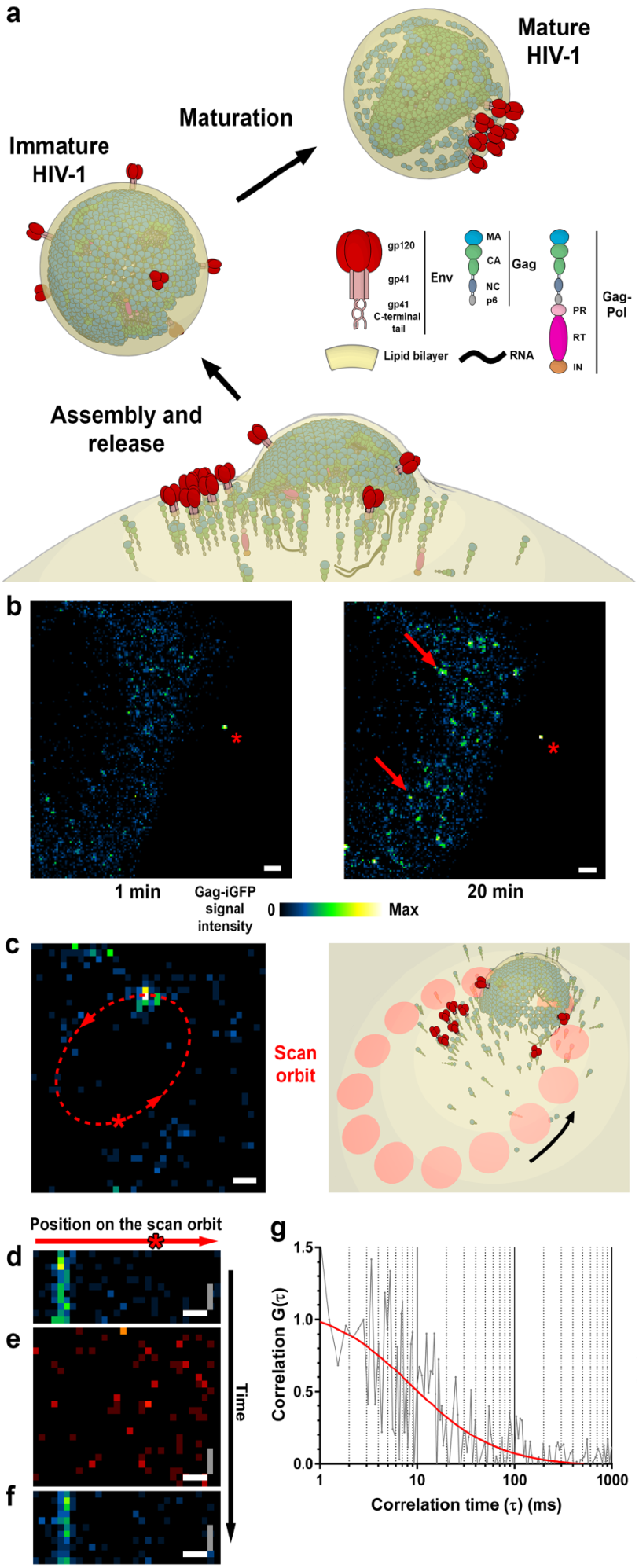
sSTED-FCS measurement of lipid diffusion inside and outside HIV-1 assembly sites. **a** Schematic illustration of Gag and Env distribution during HIV-1 assembly, release and maturation. **b** Representative confocal time-lapse images of Jurkat T-cells infected with NL4.3 Gag-iGFP HIV-1. On day 3 post infection cells were adhered to poly-L coated coverslips and monitored for the appearance of HIV-1 assembly sites prior to sSTED-FCS measurements. Images represent GagiGFP signal (blue-green-yellow) at the cell-coverslip interface at 1 minute (left) and 20 minutes (right) post adherence. Arrows indicate the appearance of the individual diffraction limited HIV-1 assembly sites, which were subsequently analysed by sSTED-FCS. A cell-free HIV-1 particle is indicated by an asterisk. Scale bar: 1 µm. **c** Left panel: live confocal imaging was used to localize HIV-1 virus assembly sites using Gag.iGFP signal (blue-green-yellow) as a guide and to align them with the position of the circular scan orbit (red dashed line). Right panel: schematic illustration of sSTED-FCS orbit over HIV-1 assembly site. Arrows indicate the direction of the scan. Scale bar: 200 nm. **d-f** Representative confocal mode signal intensity carpets for Gag.iGFP signal (blue-green-yellow) prior (**d**) and post (**f**) acquisition of STED mode intensity carpet for lipids (here phosphatidylethanolamine (PE)-ATTO 647N) (e) inside and outside HIV-1 assembly sites. Image xand y-axis correspond to the position on the scan line and signal intensity at each time point, respectively. Scale bars: x-axis (white) = 200 nm, y-axis (grey) = 1.24 ms. **g** Representative normalised autocorrelation curve of PE-ATTO 647N diffusion (grey line) obtained from a single position (marked by the red asterisk in **c** and **d-f**) on the circular scan line within correlation carpet. Autocorrelation curve (red) was fitted using generic two-dimensional diffusion model.

## Results

### PI(4,5)P2 and Cholesterol, but not sphingomyelin, are trapped during HIV-1 assembly in infected CD4+ T cells

Assembly of fully infectious HIV-1 particles requires a concomitant assembly of viral Env and Gag/GagPol at the assembly site, i.e. at the cellular plasma membrane, involving specific lipid environments that can be either pre-existing or generated by the viral proteins. To distinguish these possibilities, we infected Jurkat CD4+ T lymphocytes with infectious NL4.3 HIV-1 viruses. These viruses expressed green fluorescent protein (GFP), which was inserted between MA and CA domains of Gag (Gag-iGFP) (30). The concomitant fluorescence allowed to track the movement of fluorescently tagged lipids at HIV-1 assembly sites (Fig. 1a). 72 hours post infection we tested whether infected cells are able to support HIV1 assembly. Time lapse imaging at 37 ° C of cells adhered to poly-L-Lysine coated coverslips highlighted that infected cells produced new virus assembly sites within 20 minutes post adherence (Fig 1b) This is consistent with previous HIV1 assembly studies of HIV-1 particles (31) and HIV-1 Gag virus-like-particles production in CD4+ T cells (32). To visualize lipids mobility within or outside of virus assembly sites, we labelled the CD4+ T cells with fluorescent lipid analogues prior to adhering cells to poly-L-Lysine coated coverslips. The selected analogues correspond to lipids known to be enriched in HIV-1 membrane such as Chol and SPM, to have major role in Gag membrane-interaction such as PI(4,5)P2, or to be selectively excluded in HIV membrane such as phosphatidylethanolamine (PE). To acquire lipid mobility data via sSTED-FCS, HIV-1 assembly sites were aligned with the laser scanning orbit of our microscope, using Gag-iGFP fluorescence signal as a guide (Fig 1c). For this we first took larger confocal microscope images identifying spots of GFP fluorescence as relevant Gag clustering sites. For our analysis, we selected only small and immobile Gag clusters at the T cell surface with low fluorescence Gag signal intensity. Previous studies have shown that such sites represent virus assembly that is still in progress (32), and thus minimize the risk of performing lipid mobility analysis on the site of already budded virus particles. After this identification step, the microscope was switched from imaging (x-y axis) to continuous orbital line scanning (x-t axis, intensity carpet acquisition) mode. Consequently, we recorded temporal fluorescence intensity fluctuations at each position (or pixel) along the scanned orbit. An orbit always included Gag-clustering sites and thus HIV-1 assembly sites (GFP fluorescence) and areas without Gag (no GFP fluorescence) (Fig 1). We firstly acquired a short intensity carpet for Gag fluorescence in confocal mode to ensure that a) the scanning orbit was correctly overlapped with the position of the virus assembly site; and b) the Gag assembly site was sufficiently immobile to enable lipid mobility measurements (Fig 1d). Afterwards, we acquired an intensity carpet for the red-emission signal of the fluorescent lipid analogues with a reduced observation spot size (diameter 100 nm, STED microscopy mode) (Fig 1e). The reduced observation spot size ensures that we are indeed mainly covering lipid data at the assembly site only, and not averaging over a larger non-relevant area. Finally, we again recorded a fluorescence intensity carpet of Gag fluorescence in confocal mode to further ensure that the virus assembly site shows no detectable drift during the whole acquisition process (Fig 1f). The acquisition parameters for the recording of the fluorescent lipid analogue data were chosen for reducing bias and for generating accurate data: 1) The spot size of 100 nm in diameter ensured sufficient spatial resolution for distinguishing trapped from non-trapped lipids (see results further on) and yet still high enough signal-to-noise ratios for reasonable and reproducible FCS data (see error bars further on). 2) Orbital beam-scanning with 3.23 kHz realized a low enough dwelling at each scanned point to minimize (excitation and STED) laser exposure and thus phototoxic effects as well as a high enough temporal resolution of the decaying FCS data (34). 3) The number of total points (30) per orbital scan facilitated the collection of sufficiently enough data points for an accurate identification of the virus assembly sites (Fig. S1) (11). FCS data were generated for each point along the orbital scan and fitted with a generic twodimensional (2D) diffusion model to obtain the average transit times through the observation spot and the diffusion coefficient of the lipid analogues at each location (Fig 1g). Figure 2 shows the change in lipid lateral mobility observed in infected Jurkat T cells within (red boxes) and outside of (green boxes) the HIV-1 assembly sites. The lateral mobility of the lipids as measured in uninfected control cells are presented in blue. As previously observed for PI(4,5)P2 in SLBs ((7) and Fig. S2), the median diffusion coefficient of PI(4,5)P2 (Fig 2a) within the assembly sites was strongly reduced (D = 0.02 *µ*m^2^/s) compared to the one observed outside of the assembly sites (D = 0.17 *µ*m^2^/s) or in HIV negative cells (D = 0.20 *µ*m^2^/s). Therefore, in infected Jurkat T cells, general mobility of PI(4,5)P2 was 8to 10-fold decreased within the assembly sites compared to elsewhere or in HIV negative cells. We further tested Gag specific mobility of Chol. Chol has been shown to play a major role in the assembly process of HIV-1 virus(23). When monitoring the diffusion coefficient of Chol (Fig 2b), we observed a similar decrease of Chol mobility (D = 0.10 *µ*m^2^/s) within the assembly sites compared to elsewhere (D = 0.25 *µ*m^2^/s) and in HIV negative cells (D = 0.29 *µ*m^2^/s). In contrast, SPM did not exhibit any significant change in mobility in the presence or absence of HIV-1 or within and outside the assembly sites (Fig 2c; D = 0.15 and 0.17 *µ*m^2^/s within and outside the assembly sites, respectively, and D = 0.15 *µ*m^2^/s in uninfected cells) Similarly, we observed no significant change in the mobility of the PE lipid analogue within (D = 0.32 *µ*m^2^/s) versus outside (D = 0.31 *µ*m^2^/s) the assembly sites or versus uninfected cells (D = 0.31 *µ*m^2^/s) (Fig 2d).

**Fig. 2.**
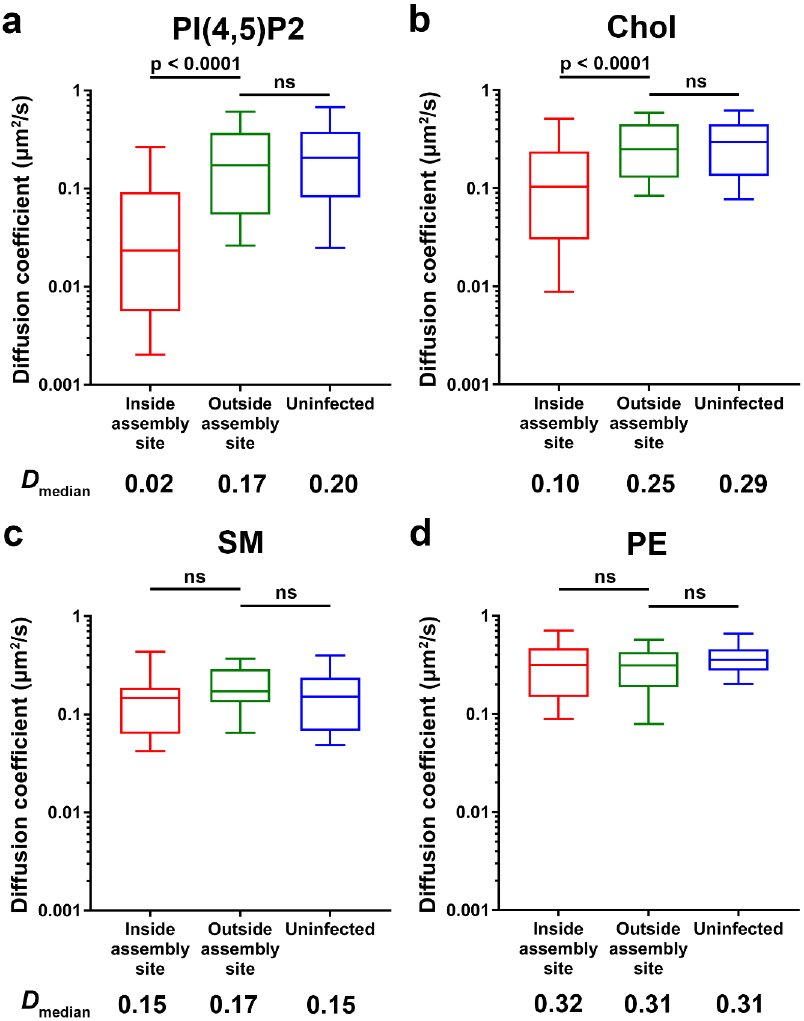
Lipid mobility changes in NL4.3 Gag-iGFP HIV-1 infected Jurkat T-cells. Median lipid diffusion coefficients (D) as determined by sSTED-FCS measurements of 20 sites each from two independent virus preparations and infections. Box and whisker plots (horizontal line - median, box 25-75 % percentiles or interquartile range (IQR) and whiskers - 10 90 % measurements) showing D values inside assembly sites (red), outside assembly sites (green) and in non-infected control cells (blue). Data was collected for phosphatidylinositol 4,5-bisphosphate (PI(4,5)P2) (**a**), cholesterol (Chol) (**b**), sphingomyelin (SPM) (**c**), and phosphatidylethanolamine (PE) (**d**). Statistical significance was assessed by Wilcoxon rank-sum test.

### Expression of HIV-1 Gag alone is sufficient to immobilize PI(4,5)P2 and Cholesterol during formation of Virus Like Particles

HIV-1 Gag alone is able to self-assemble at the plasma membrane of the Jurkat T cells, leading to the generation of immature virus-like-particle (VLP). Kerviel et al. (35) proposed that self-association of Gag is sufficient to generate a specific lipid environment. Therefore, by transfecting HIV-1 Gag only expression vector into Jurkat T cells and measuring changes in the mobility of lipids at Gag self-association sites, we tested whether the selective lipid clustering we have observed in HIV-1 infected cells (see above) were also present in Gag only assembly sites. As with the infected Jurkat T cells, we have firstly tested for the evidence of appearance of VLP assembly sites at the surface of HIV Gag expressing Jurkat cells (Fig 3a), followed by the determination of lipid diffusion coefficients both within and outside of those sites. We determined lipid diffusion at arbitrary sites in non-transfected (non-Gag expressing) cells as control. Figure 3b-e depicts the different values of diffusion coefficients determined within (red boxes) and outside of the assembly sites of HIV Gag VLP (green boxes), and in the non-transfected control cells (blue box). As observed in infected cells, the lateral mobility of PI(4,5)P2 was significantly reduced inside (D = 0.11 *µ*m^2^/s) compared to outside of assembly sites or to non-transfected cells (D = 0.24 *µ*m^2^/s and 0.20 *µ*m^2^/s, respectively) (Fig 3b). However, the reduction in mobility between inside and outside the assembly sites is only 2.2-fold (0.24/0.11) for HIV Gag VLP only expressing T cells in comparison with 8.5-fold (0.17/0.02) for HIV infected T cells. Nevertheless, experiments on model membranes support the direct role of Gag in this process of PI(4,5)P2 trapping. Upon addition of Gag to supported lipid bilayers (SLBs) with 0.2% of fluorescent PI(4,5)P2 we observed the formation of PI(4,5)P2 clusters at the Gag binding sites with a concomitant 7-fold decrease in PI(4,5)P2 mobility at these sites compared to elsewhere on the SLBs or on SLBs in the absence of Gag (Fig. S2). Also in line with our data on infected T cells, we also observed a 2-fold reduction in the lateral mobility for Chol within the Gag assembly sites (D = 0.16 *µ*m^2^/s) of HIV Gag VLP only expressing T cells, compared to either outside of assembly site (D = 0.30 *µ*m^2^/s) or in non-transfected control cells (D = 0.29 *µ*m^2^/s, Fig 3c). In contrast, SPM and PE again exhibited no significant changes in their lateral mobility between HIV-1 Gag self-assembly sites when compared to outside the sites or in non-transfected cells (Fig 3d and Fig 3e respectively).

**Fig. 3.**
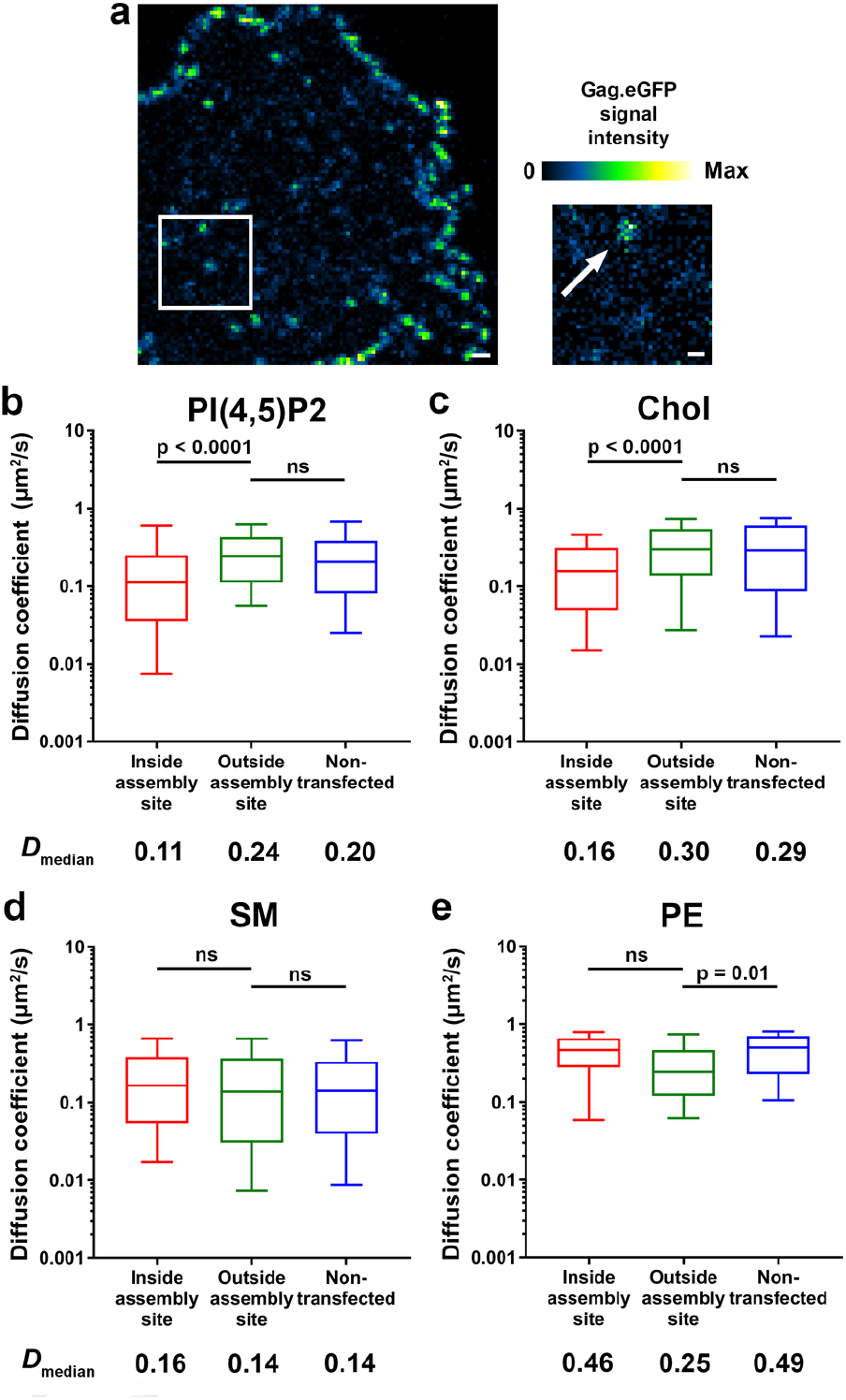
Lipid mobility changes in HIV-1 Gag.iGFP transfected Jurkat T cells. a Representative live confocal image of Jurkat T cells expressing HIV-1 Gag.iGFP. Image represents Gag.iGFP signal (blue-green-yellow) at the cell-coverslip interface at 24 hours post transfection. Scale bar: 1 µm. Zoomed in confocal image depicts an individual HIV-1 assembly site indicated by an arrow. Scale bar: 200 nm. b-e Median values of the lipid diffusion coefficients (D) were determined for phosphatidylinositol 4,5-bisphosphate (PI(4,5)P2) (**a**), cholesterol (Chol) (**b**), sphingomyelin (SM) (**c**), and phosphatidylethanolamine (PE) (**d**) by sSTED-FCS on 20 sites each from two independent Gag.iGFP transfections. Box and whisker plots (horizontal line - median, box 25-75 % percentiles or interquartile range (IQR) and whiskers - 10 90 % measurements) showing D values inside assembly sites (red), outside assembly sites (green) and in untransfected control cells (blue). Statistical significance was assessed by Wilcoxon rank-sum test.

### HIV-1 Gag is the primary determinant of lipid trapping efficiency at HIV-1 assembly sites in CD4+ T cells

The decrease in lateral mobility of PI(4,5)P2 and Chol at the Gag assembly sites suggests trapping of these lipids during virus particle formation. To quantify this restricted diffusion further, we utilized a confinement index based on normalized relative cumulative frequency histograms of the values of the diffusion coefficient D (for D = 0.001 to 1 *µ*m^2^/s, normalized to 100 for D = 1 *µ*m^2^/s). Representative cumulative frequency histograms for diffusion within and outside of the assembly sites are shown in Figure 4a for the fluorescent PI(4,5)P2 lipid analogue (and in Fig. S3, S4 for other lipid analogues), highlighting a huge difference for PI(4,5)P2 and Chol (but not for SPM and PE). To specify this difference further, we classified three diffusion regimes, “fast”, “intermediate”, and “slow” mobility, for respective values of D ≤ 0.01 *µ*m^2^/s, 0.01 D ≥ 0.1 *µ*m^2^/s, and D ≥ 0.1 *µ*m^2^/s, we calculated the sum of cumulative frequencies within each regime, and plotted the confinement index for each regime as ratio of the latter values for lipid diffusion within and outside of the assembly sites, or within the assembly sites and non-infected (or non-transfected) cells. For an increased confinement of lipids inside the assembly sites, we would expect a larger relative fraction of low values of D, i.e. an increased fraction and thus confinement indices > 1 within the “intermediate” and especially the “slow” mobility regime (and consequently a reduced fraction and thus a confinement index < 1 within the “fast” regime). In contrast, similar cumulative frequency histograms and thus confinement indices 1 for all regimes indicate no change in mobility and thus no confinement within the assembly sites. As expected, the various values of confinement indices confirm our previous conclusion, i.e. that SPM and PE did not exhibit any confinement within assembly sites, while PI(4,5)P2 and Chol revealed a drastic reduction of lateral mobility and thus confinement at the assembly sites. When we compared the confinement index values obtained for the different lipids in HIV-1 Gag transfected cells (Fig 4b, grey bars) against those in HIV-1 infected cells (Fig 4b, black bars), we observed similarities between these two conditions. Specifically, both PI(4,5)P2 and Chol have values of high confinement indices > 1 in the “intermediated” and “slow” regime for both HIV infected and Gag-only expressing T cells, while values 1 of confinement indices are conserved for both SPM and PE. This highlights that the Gag protein on its own seems to be sufficient to facilitate the confinement or trapping of cholesterol and PI(4,5)P2 in HIV assembly sites, and no other molecular factor only present in infected cells (and not in the Gag-only case) is required.

**Fig. 4.**
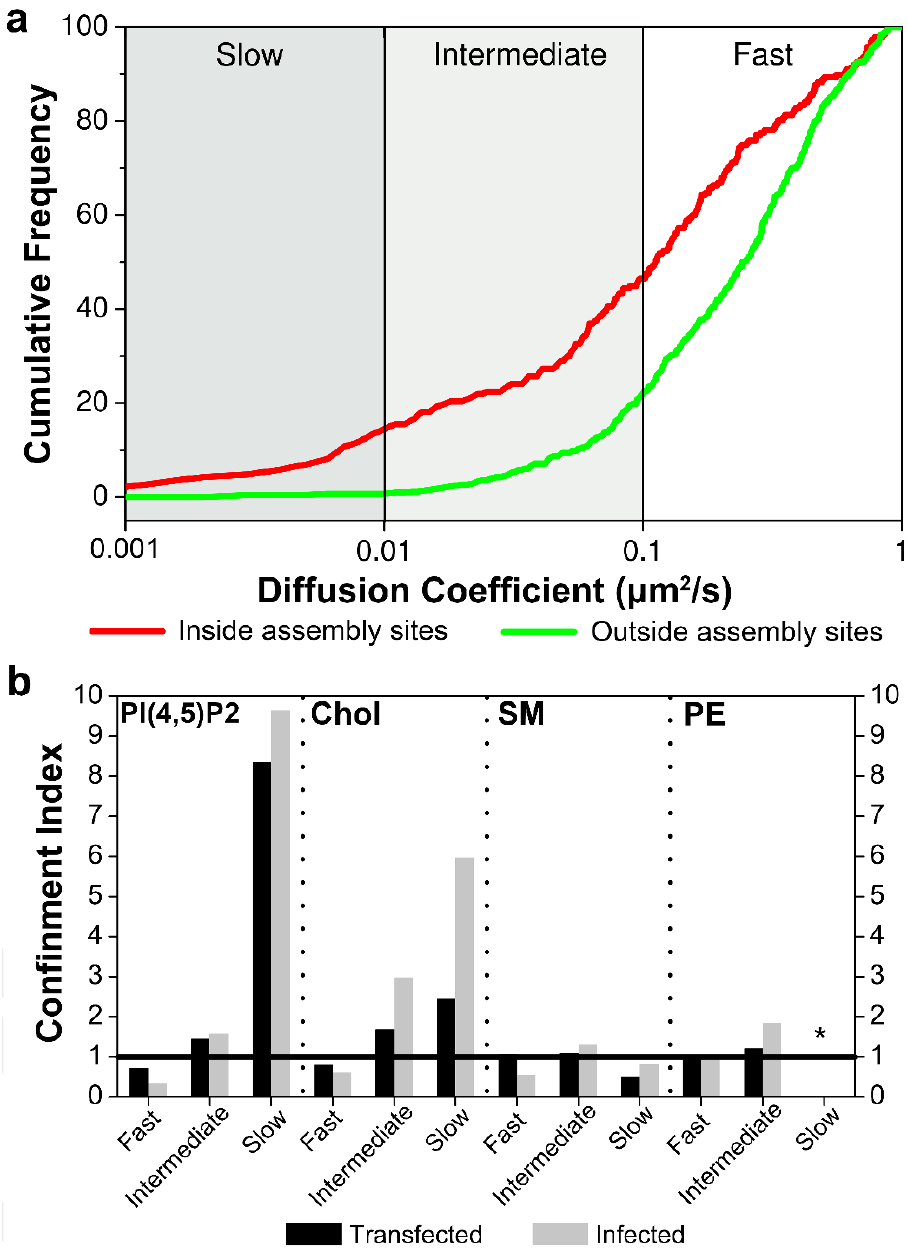
Lipid trapping induced by Gag during HIV-1 assembly in infected or transfected Jurkat T cells. Enrichment indices based on the relative normalized cumulative frequency distribution of the diffusion coefficients D (for D = 0.001 to 1 *µ*m^2^/s, normalized to 100 for D = 1 *µ*m^2^/s), which were arbitrarily divided into three distinct diffusion regimes, “slow” (D < 0.01 *µ*m^2^/s), “intermediate” (0.01 ≤ D < 0.1 *µ*m^2^/s) and “fast” (D ≥ 0.1 *µ*m^2^/s). The sum of cumulative frequencies within each regime resulted in values of the confinement index for each regime as ratio of the latter values for lipid diffusion within and outside of the assembly sites, or within the assembly sites and non-infected (or non-transfected) cell. **a** Cumulative distribution of diffusion coefficients obtained for PI(4,5)P2 in HIV-1 Gag transfected cells inside the assembly site (red line) and outside the assembly site (green line, see supplementary data 3 and 4 for cumulative distributions of the other lipids). **b** Values of the confinement indices found for PI(4,5)P2, Chol, SM and PE for HIV-1 Gag transfected (grey bars) and HIV-1 infected (black bars) Jurkat T cells. Chol and PI(4,5)P2 show strong confinement (indices > 1 for “intermediate” and especially “slow” regimes) inside assembly sites for both transfected only or fully infected Jurkat T cells, while neither SPM nor PE exhibited confinement inside assembly sites (confinement index 1 for all regimes). *: No diffusion coefficients below 0.01 *µ*m^2^/s were observed for PE.

## Discussion

How enveloped viruses acquire their lipid membrane shell is of great importance for their biogenesis and their capacity to infect cells. In the case of HIV-1, recently it has been shown that the envelope lipid composition was a determinant of low molecular mobility of the Env protein (3, 13). Several works have been conducted to reveal different envelope viruses having their own unique lipid composition (2, 4, 6, 36), implying their distinct biological requirements during infection, and suggesting their capacity in sorting specific host lipids into their virus membrane. Two main lipids and a sterol have been described to play a major role for HIV-1 biogenesis and infectivity. PI(4,5)P2 is known to be the plasma membrane lipid targeted by the MA domain of the polyprotein Gag (20, 21, 27, 37). Sphingomyelin and Cholesterol enriched lipid environments, often denoted “raft” like domains, have also been described as essential for envelop retrieving (38), HIV-1 assembly (23, 39, 40) or infectivity (24, 25). However, it is unknown whether a pre-existing cellular lipid complex or a virus induced lipid platform is the source of virion lipid membrane during HIV-1 formation (35). Lipid sorting and lipid domain generation in viral assembly can be identified by nanoscale spatial changes in lipid mobility at the plasma membrane (10, 11), and our super-resolution microscopy based approaches for the analysis of the lipid dynamics in living cells represent the most relevant model of observation. In this study, we have determined the diffusion coefficients of lipids by applying scanning STED-FCS (sSTED-FCS) on HIV infected, Gag expressing, and HIV negative CD4+ Jurkat T cells. sSTED-FCS approach allows for direct simultaneously quantitative comparison of lipids mobility both within or outside assembly sites at the plasma membrane. In comparison with the plasma membrane of HIV-1 negative cells or plasma membrane regions outside of HIV-1 assembly site, the mobility of PI(4,5)P2 and Chol within viral assembly sites is drastically reduced (up to 10 times decrease of median D). Interestingly, the same trend was observed on the assembly site of Gag VLPs in cells expressing only Gag. Indeed, although to a less extent than in HIV-1 producing cells, the median values of the diffusion coefficients of PI(4,5)P2 and Chol were significantly reduced in the VLP assembling area. In contrast, the mobility of neither SPM nor neutral lipid PE exhibited any variation in median values, for both HIV-1 producing or Gag VLPs producing cells. It has been recently shown that diffusion coefficient distributions could unravel the nature of lipid motions (41). Therefore, to allow direct comparison of the virus component effect on lipid mobility, independently of the cell or the number of expressed component (Gag alone or infectious virus), a confinement index was defined. As in more detail described within the result section, it is based on the relative changes in three different parts of the normalized cumulative distribution of the logarithm of the diffusion coefficients observed inside and outside the virus or VLP assembling site. The value of this confinement index into these three parts reflects the relative increase in lipids that are trapped (“slow”, D < 0.01 *µ*m^2^/s), confined (“intermediate”, 0.01 < D < 0.1 *µ*m^2^/s) or freely diffusing (“fast”, D > 0.1 *µ*m^2^/s) at the assembly sites. By definition, the confinement index will be equal to 1 if no effect occurs on lipid mobility inside the assembly when compared to outside the assembly site. This unit value was always observed with the PE lipid which is generally used as a control of free diffusing lipid in cell plasma membranes (10) and is not described as being enriched in HIV-1 envelop (1, 4). Amongst the different lipids tested here, the PI(4,5)P2 exhibited the strongest confinement index for the slow part of the distribution, (c.i. 10) both for HIV-1 virus and VLP assembly sites in CD4+ T cells. This reveals that transient binding of PI(4,5)P2 to aggregating Gag conjugated PI(4,5)P2 into the nascent virion or the VLP. This assembly site specific trapping of PI(4,5)P2 seems to be indeed exclusive to Gag since the observed confinement indices showed the same trend independent of assembling HIV-1 or VLPs. Moreover, similar confinement indices were observed on the SLB in the in vitro system tested here, i.e. SLBs made of PC, PS and fluorescent PI(4,5)P2 with or without addition of self-assembling HIV-1 Gag protein (see Supplementary Data 2). Similar to PI(4,5)P2, the confinement index of slow diffusing Chol was also above 1 in infected cells as well as in transfected cells, confirming that Chol is trapped during HIV-1 assembly and that Gag alone is sufficient to restrict cholesterol mobility at the assembly site. Consequently, restriction of PI(4,5)P2 and Chol seems to be independent of non-Gag viral proteins, such as HIV-1 Env or accessory/regulatory proteins. Using in vitro lipid membrane models of different lipid compositions, we recently reported (7) that Gag was able to generate PI(4,5)P2-cholesterol but not SPM enriched lipid nanodomains. This is consistent with data observed here in HIV1 producing or Gag expressing CD4+ T cells. Notably, such local enrichment of PI(4,5)P2 and Chol lipids at virus assembly sites has been predicted using coarse grain molecular dynamics in the case of the matrix self-assembly of Ebola virus (42), implying a potential general mechanism for a number of enveloped viruses that bud from the host cell plasma membrane. Enrichment of specific lipid in HIV-1 viruses has already been observed (2, 4, 23) but the mechanism remains unknown. Different values of virion lipid enrichment have been found, depending on the cell lines and the biochemical methods to purify viruses or isolate plasma membranes from the other cell membranes (1, 2, 4). In H9 T lymphocytes, Chan et al. have shown that HIV-1 lipid envelop is two times higher in PI(4,5)P2 and three times higher in cholesterol when compared to the plasma membrane of the HIV-1 producing H9 cells. Our results show that this enrichment occurs during virus assembly. Given that PI(4,5)P2 is important for the binding of Gag to the cell plasma membrane (28), the clustering of PI(4,5)P2 at the cell membrane may provide an additional mechanism to further concentrate Gag thereby enhancing virus assembly. Furthermore, molecular simulations of PI(4,5)P2 clustering has revealed the induction of negative membrane curvature (43) by PI(4,5)P2, which could facilitate the formation of spherical particle structure during virus assembly. Cholesterol has been shown to play a role in HIV-1 Gag matrix binding to membrane(5, 44) as well as in the stabilization of PI(4,5)P2 lipid nanodomains in the absence of proteins (43). Therefore, it would be difficult to discriminate whether cholesterol trapping is a consequence of PI(4,5)P2 membrane enrichment at HIV-1 assembly sites or is due to a direct interaction with Gag. Nevertheless, the cholesterol confinement index found in this work is much lower than the one observed for PI(4,5)P2, allowing the conclusion that the incorporation of cholesterol at the HIV-1 assembly site is more of a consequence of PI(4,5)P2 enriched membrane environment than a strong direct interaction with Gag. It has been suggested for a long time that pre-existing sphingomyelin-cholesterol enriched cell plasma membrane nanodomains could play a role in HIV-1 assembly (reviewed in (38)). Therefore, changes in SPM dynamics at the virus assembly site were also assessed using our sSTED-FCS approach. The observed confinement indices of SPM were never above 1, highlighting that HIV-1 assembly has no effect on the SPM mobility at the CD4+ T cell plasma membrane. Overall our results provide direct evidence that HIV1 Gag assembly drives the lipid sorting in host CD4 T cell plasma membrane and creates its own PI(4,5)P2/cholesterol lipid enriched membrane environment likely to provide a positive feedback loop to recruit additional Gag proteins at the budding site through PI(4,5)P2-Gag interactions for the biogenesis of HIV-1 particles.

## Material and Methods

### Experimental Design

The main goal of this study was to unravel whether the viral Gag protein is targeted towards preexisting lipid domains at the host cell plasma membrane that will help in HIV-1 assembly, or whether, in the opposite, Gag creates its own lipid domain during virus formation. In order to do so, we measured the change in mobility of different lipids at HIV-1 assembly sites. We used lipids known to play a role either in Gag membrane targeting (PI(4,5)P2) and assembly (Cholesterol) or known to be part of pre-existing lipid domains at the plasma membrane (SM) and finally a control lipid (PE) not described to play a role in any of the two possible mechanisms. To achieve our goal, our experimental setup was based on a direct measurement of apparent diffusion coefficient of the lipids inside and outside HIV-1 Gag assembly sites using spatial scanning fluorescence correlation spectroscopy on a super-resolution STED microscope (delivering an observation spot size of around 100 nm in diameter, i.e. below that of a conventional microscope). Our measurements were performed in HIV-1 infected CD4 T-cells and as a control in CD4 T-cells expressing Gag only. Finally, using a top-bottom approach of purified HIV-1 Gag on PI(4,5)P2supported lipid bilayers we could highlight that Gag was sufficient to generate a PI(4,5)P2-nanodomain platform for assembly.

### Lipids and fluorescent lipid analogues

Lipids 1,2dioleoyl-sn-glycero-3-phosphocholine (DOPC), brain L-αphosphatidylserine (PS) and brain Phosphatidylinositol 4,5biphosphate (PI(4,5)P2) were purchased from Avanti Polar lipids (Alabaster, AL, USA). ATTO 647N – 1,2-dipalmitoylsn-glycero-3-phosphoethanolamine (ATTO647N-DPPE) and ATTO 647N – Sphingomyelin (ATTO647N-SM) were purchased from Atto-Tec GmbH (Siegen-Weidenau, Siegen, Germany). Abberior STAR RED (KK114) 1,2-dihexadecanoyl-sn-glycero-3-phosphoethanolamine (KK114-DPPE), Cholesterol – PEG(1k) Abberior STAR RED (KK114) (Chol-PEG(1k)-KK114), and ATTO 647N phosphatidylinositol 4,5-biphosphate (ATTO647NPI(4,5)P2) were all purchased from Abberior GmbH (Göttingen, Germany).

### Gag protein and Supported Lipid Bilayers

Full length Gag protein has been produced and purified as described previously(45). Supported Lipid Bilayers (SLBs) for Gag lipid clustering measurements were prepared from 30nm small unilamellar vesicles (SUVs) with DOPC - 68%, PS - 30%, PI(4,5)P2 - 1.9% and ATTO647N-PI(4,5)P2 0.1% mol at 0.1 mg/mL in a citrate buffer (Na Citrate 10mM, 100 mM NaCl, and 0.5 mM EGTA, pH 4.6). The SUV were then spread for 45 min at 37°C on coverslips pre-treated by sonication in 5% SDS solution and incubation for 30 min in freshly made pi-ranha solution (H2SO4:H202 2:1 vol). After disposition of the SLBs scanning FCS measurements were performed and again 15 minutes after Gag was added at 1*µ*M final concentration.

### Plasmids

Plasmid expressing fully infectious T cell tropic NL4.3 HIV-1 Gag-iGFP was previously described (30) and was a kind gift from Dr Benjamin Chen. Plasmid expressing NL4.3 HIV 1 Gag eGFP fusion protein was previously described (45) and was a kind gift from Dr Hugues de Rocquigny.

### Cell culture

Jurkat T cells (Human T cell leukemia cell line– ATCC TIB-152) were grown in RPMI-1640 with Glutamax (GIBCO), supplemented with 10 % foetal calf serum and, 100 U/mL penicillin-streptomycin and 20mM HEPES pH 7.4. Cells were maintained at 37 °C, 5 % CO2.

### NL4.3 HIV-1 Gag.iGFP particle preparation

Fully infectious NL4.3 HIV-1 Gag-iGFP particles were prepared from the tissue culture supernatant of 293T cells transfected using polyethyleneimine (PEI) with 15 *µ*g pNL4.3 HIV-1 Gag-iGFP plasmid. Tissue culture supernatants were harvested 48 h after transfection, and particles were concentrated using Lenti-X Concentrator reagent (Clontech) according to the manufacturer’s instructions. Concentrated particles were snap frozen and stored in aliquots at −80 ° C

### Jurkat T cell infection

Jurkat CD4+ T cells were infected by incubation of 1 million cells with HIV-1 NL4.3 Gag-iGFP particles in 50 uL RPMI medium for 1 h at 37 °C. Cells were washed three times in RPMI medium and then cultured for 72 h at 2 million cells/mL to achieve 5-10 % infection rate with progeny virus production.

### Jurkat T cell transfection

Jurkat T cells (2×106) were microporated with 4 *µ*g of plasmid expressing NL4.3 HIV-1 Gag eGFP, using the AMAXA system (Lonza), as previously described (46). After microporation, Jurkat T cells were incubated in RPMI complete medium, and harvested 24 h posttransfection for microscopy acquisitions as described in the following sections.

### Microscope setup

Scanning STED-FCS measurements were performed on the Abberior Instrument Expert Line STED super-resolution microscope (Abberior Instruments GmbH, Göttingen, Germany) using 485 nm and 640 nm pulsed excitation laser sources and a pulsed STED laser operating at 775 nm and 80 MHz repetition rate. The fluorescence excitation and collection was performed using a 100×/1.40 NA UPlanSApo oil immersion objective (Olympus Industrial, Southend-on-Sea, UK). All acquisition operations were controlled by Imspector software (Abberior Instruments) and point FCS data for the calibration of the observation spot sizes was recorded using a hardware correlator (Flex02-08D, correlator.com, operated by the company’s software). Confocal microscope FCS measurements were performed on a Zeiss LSM 780 (Zeiss, Iena, Germany) using a HeNe 633 nm laser as the excitation source. Fluorescence excitation and collection was performed using a 63×/1.40NA PlanApo oil immersion objective (Zeiss). Acquisition operations were controlled by the Zeiss Zen software. Point FCS was used to calibrate the observation spot with the LSM 780 internal hardware correlator.

### Microscope sample preparation

For calibration of the observation spot sizes using STED-FCS, supported lipid bilayers (SLBs) were prepared, as previously described (47), by spin coating a coverslip with a solution of 1 mg/mL DOPC and 0.5 μg/mL KK114-DPPE in CHCl3/MeOH. Coverslips were cleaned by piranha solution (3:1 sulfuric acid and hydrogen peroxide). The lipid bilayer was formed by rehydrating with SLB buffer containing 10 mM HEPES and 150 mM NaCl pH 7.4. Calibration of the confocal observation spot size was made with a 50 nM solution of Rhodamine in pure water using D=280*µ*m^2^.s^-1^ as the diffusion coefficient. For lipid mobility measurements inside and outside of Gag assembly sites in mammalian cells 3×105 Jurkat T cells 72 h post-infection with NL4.3 HIV-1 Gag-iGFP virus particles or 24 h post-transfection with NL4.3 HIV-1 Gag-eGFP expressing plasmid were resuspended in 200 *µ*L of Leibovitz’s L 15 medium (Sigma) and incubated with 25 nM of the indicated fluorescent lipid analogue probe for 5 min at 37 °C. Cells were then washed three times in L-15 medium, resuspended in 250 *µ*L L-15 medium and adhered to poly-Llysine (Sigma) coated glass surface of 8-well Ibidi *µ*-Slides (Ibidi, Martinsried, Germany) at 37 °C prior to microscopy measurements. For lipid mobility measurements on Myr(-) Gag containing SLBs, prior to experiments citrate buffer was changed to protein buffer (Hepes 10 mM, pH=7.4, KCl 150 mM, EDTA 2mM), followed by injection of 1 *µ*M Myr(-)Gag.

### Scanning (STED)-FCS signal acquisition

sSTED-FCS data of the lipid mobility inside and outside Gag assembly sites in Jurkat T cells was acquired at 37 °C using fluorescent Gag signal as a guide. Fluorescent lipid signal intensity fluctuation carpets (scanning an orbit multiple times) were recorded using the Imspector software with following parameters: orbital scan frequency 3.23 kHz, scan orbit length 1.5 *µ*m, pixel dwell time 10 *µ*s, total measurement time 10 s, pixel size 50 nm/pixel, 10 *µ*W excitation power (back aperture) at 640 nm, observation spot diameter 100 nm FWHM (as determined by SLB calibration measurements – Supplementary Data 1). The scan frequency selected for these experiments was set sufficiently high in order to enable the extraction of the diffusion dynamics of fluorescent lipid analogues in the cell plasma membrane (11).

### FCS curve autocorrelation and fitting

The FoCuS-scan software(34) was used to autocorrelate the scanning STEDFCS data, and to fit it with a classical 2D diffusion model:

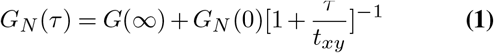

where *G*_N_(τ) is the correlation function at time lag τ *G*_N_(∞) the offset, *G*_N_(0) the amplitude, *t_xy_* the average lateral transit time through the observation spot. As done in all previous scanning STED-FCS measurements, we left out anomaly factors in the fitting equation (as often done in non-scanning single point STED-FCS data), since the decay of the scanning STED-FCS curves starts at 0.5-1 ms correlation times only and because anomaly is much better revealed by comparing (cumulative) frequency histograms of resulting parameters(11). Values of the molecular diffusion coefficient (D) were calculated from txy:

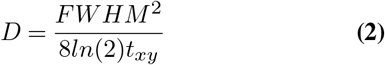

where FWHM represents the usual full-width half maximum of the observation spot (100 nm, see Supplementary Data 1).

### Calculation of the confinement index

Diffusion coefficients obtained for the different lipid in the different conditions tested were binned in groups every 10^−3^*µ*m^2^/s from 0 to 1.2 *µ*m^2^/s. The distribution was cumulated and normalized. It was then split into three parts: Slow (D < 10^−2^*µ*m^2^/s), intermediate (10^−2^ ≤ D < 10-1 *µ*m^2^/s) and fast (D ≥ 10^−1^*µ*m^2^/s). The relative proportion (Pi) of the total distribution (0 ≤ Pi ≤ 1) of each of these parts were then calculated for each molecule inside assembling site, outside assembling site and in non-infected nor transfected cells. The confinement index is defined for each of the three parts as the quotient of the Pi value found inside the assembly site over the Pi value found outside the assembly site or in non-infected nor transfected cells. (Fig 4).

### Statistical analysis

Due to the non-Gaussian nature of the diffusion coefficient data distribution, Wilcoxon test was used with the result p < 0.05 considered statistically significant. Statistical tests were performed using Graphpad Prism software. Power calculations confirmed that for chosen sample sizes the power of a two-sided hypothesis test at the 0.05 significance was over 90%. In all compared groups interquartile range (IQR) serves as an estimate of variation. IQRs were similar in all compared groups.

## ACKNOWLEDGEMENTS

We would like to thank Thomas J Hope (Northwestern University, Chicago) and the Department of NanoBiophotonics (Max Planck Institute for Biophysical Chemistry, Gottingen, Germany) for the eGFP.Vpr construct. We would also like to thank the Montpellier RIO Imaging facilities (MRI, Montpellier France) and the Wolfson Imaging Centre Oxford (UK), for help with the use of microscopes. Cyril Favard and Delphine Muriaux are members of the CNRS consortium ImaBIO, France. The overall project was funded by CNRS and the French National Agency for Research (ANR “FLUOBUDS”, ANR-13-BSV5-0006-01) and the French Agency against AIDS (ANRS). JC and CE were supported by the Medical Research Council (grant number MC-UU-12010/unit programs G0902418 and MCUU-12025), MRC/BBSRC/EPSRC (grant MR/K01577X/1), Wellcome Trust (grant 104924/14/Z/14), Deutsche Forschungsgemeinschaft (Research unit 1905 “Structure and function of the peroxisomal translocon”), and Oxford internal funds (EPA Cephalosporin Fund and John Fell Fund). Johnson Mak is recipient of funding from Australian National Health and Medical Research Council project grants App543107 and App1121697, and is a sub-awardee of National Institute of Health NIH RO1 GM064347. CF, JC, CE and DM designed the study. DM, CF, JC, PM and NY performed the experiments. JM provided the recombinant myr(-)Gag protein. CF developed the confinement index analysis. CF, JC and CE analysed the STED-FCS data. JC, CF, DM and CE wrote the manuscript. All authors contributed in discussing the data, experiments and the manuscript.

## Supplementary data

### Supplementary Note 1: STED microscope calibrationg

The diameters of the observation spots employed in the scanning STED-FCS was estimated at 37 °C following a standard calibration procedure on supported lipid bilayers (SLBs)(48), comparing transit times of freely diffusing fluorescent lipids (KK114-DPPE) in confocal and STED microscopy modes (Supplementary Fig. 1).

**Fig. S1.**
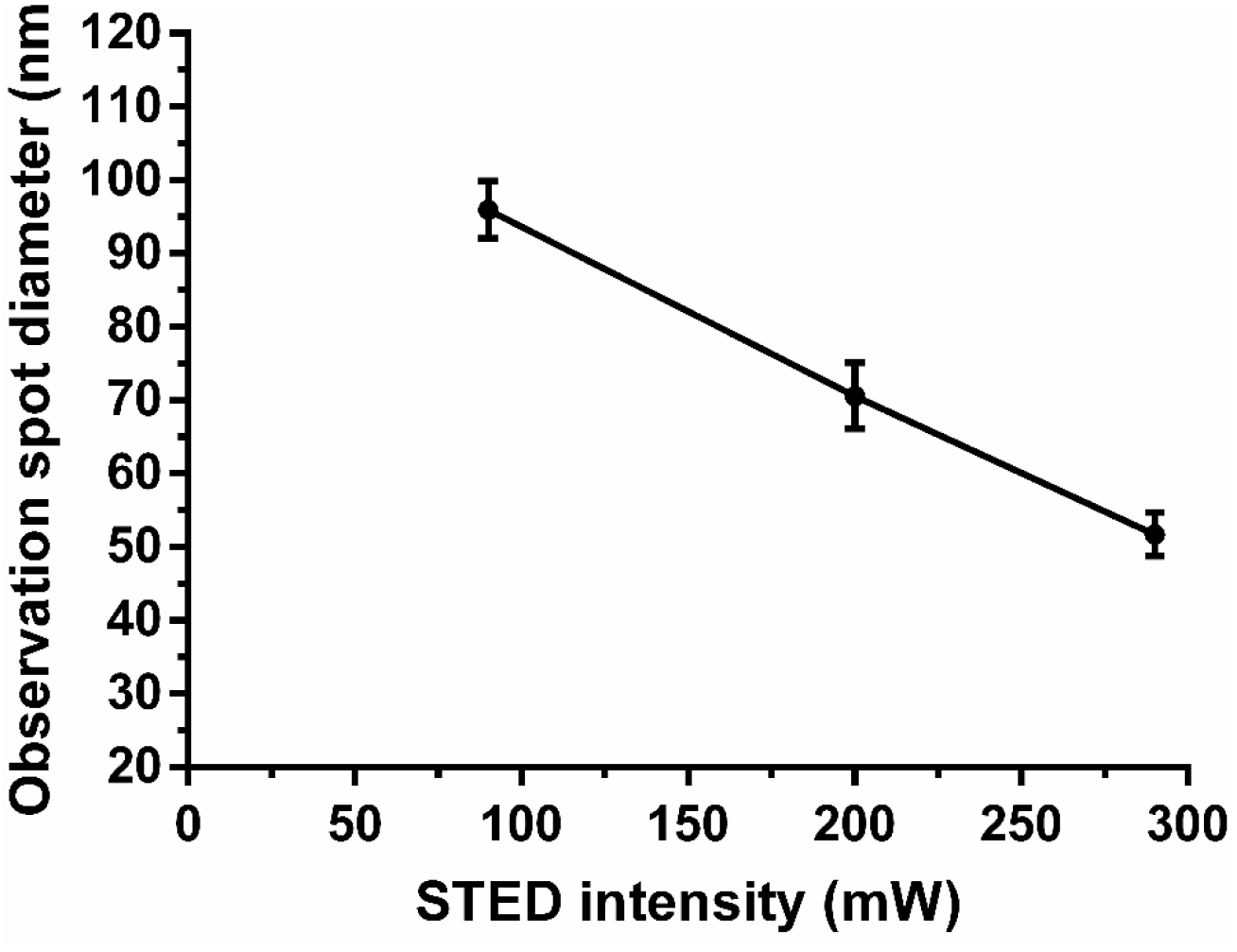
STED microscope calibration results. (**a**) Mean ± s.d. of observation spot diameters at different STED laser powers was determined by STED-FCS calibration measurements of KK114-DPPE diffusion in supported lipid bilayers (SLBs). Results represent 10 measurements each from 2 separate preparations. For the virus assembly sites measurements, a minimum STED laser power (90 mW) was set to generate an observation spot diameter (100 nm) below the diameter of HIV-1 particles (∼140 nm diameter) and virus assembly sites, while maintaining as high as possible signal-to-noise ratio and minimising the laser light exposure of live Jurkat T cells.

### Supplementary Note 2: PI(4,5)P2 clustering and trapping by full length HIV-1 Gag on biomimetic membranes

Yandrapalli et al.(7) previously reported the ability of purified full length Gag to segregate PI(4,5)P2 and cholesterol while self-assembling on supported lipid bilayers (SLB). Here, we checked the ability of Gag to induce PI(4,5)P2 clustering and trapping on SLBs with the basic lipid composition (see supplementary methods) described in Yandrapalli et al.(7) by imaging the change in Atto-647N PI(4,5)P2 fluorescence before and after addition of Gag on SLBs (Supplementary Fig. 2a and Supplementary Movie 1). As previously reported, we observed the generation of Atto-647N PI(4,5)P2 enriched domains few minutes after injection of recombinant full length Gag (Supplementary Fig. 2a). Concomitantly, confocal line scanning FCS (sFCS) was performed on SLBs in order to monitor the changes in lateral PI(4,5)P2 mobility induced by Gag self-assembly. We first measured the lateral diffusion of Atto-647N PI(4,5)P2 before Gag injection (Supplementary Fig. 2b, blue box) and compared it to the lateral diffusion observed after Gag injection inside (Supplementary Fig. 2b, red box) and outside (Supplementary Fig. 2b, green box) the generated Gag aggregation sites. We observed a significant decrease in Atto-647N PI(4,5)P2 lateral mobility at (D = 0.15 *µ*m^2^/s) compared to outside (D = 1.1 *µ*m^2^/s) the sites or before the addition of Gag (D = 0.95 *µ*m^2^/s), highlighting strong trapping of PI(4,5)P2 in lipid domains generated by Gag. We then applied the confinement index calculation to determine the enrichment of PI(4,5)P2 in Gag induced clusters. We obtained values directly in line with either HIV-1 Gag transfected cells or HIV-1 infected cells. (Supplementary Fig. 2b)

**Fig. S2.**
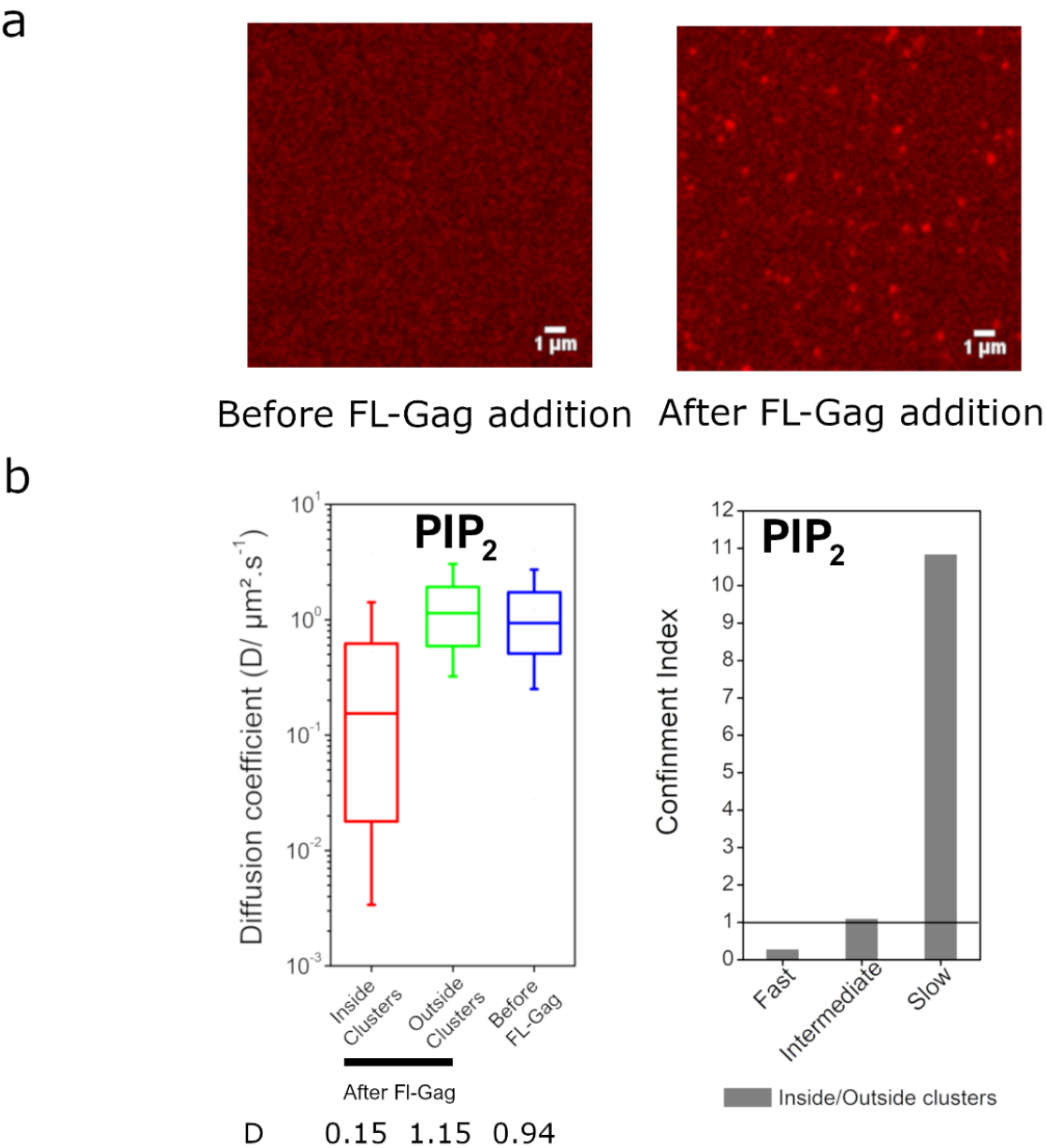
Changes in the lateral mobility of PI(4,5)P2 upon addition of Gag on supported lipid bilayers (SLB). (**a**) Confocal images of PC/PS/PI(4,5)P2 supported lipid bilayers labelled with Atto-647N PI(4,5)P2 before (upper part) and 3 minutes after addition of the full length HIV-1 Gag protein. (**b**) Box plot of the diffusion coefficients of Atto-647N PI(4,5)P2 observed before addition of Gag (blue) and 3 minutes after addition of Gag inside (red) or outside PI(4,5)P2 clusters (green), (boxes are median, first and third quartile and 10-90% whiskers). Values of the confinement indices of PI(4,5)P2 in SLB for the three different diffusion regimes (slow D < 0.1 *µ*m^2^/s, intermediates 0.1 < D < 1*µ*m^2^/s) and fast (D>1 *µ*m^2^/s). As in the case of infected and transfected cells, PI(4,5)P2 is strongly enriched in the clusters observed 3 min after addition of Gag.

### Supplementary Note 3: Cumulative frequency distributions observed in infected cells for the different lipids

**Fig. S3.**
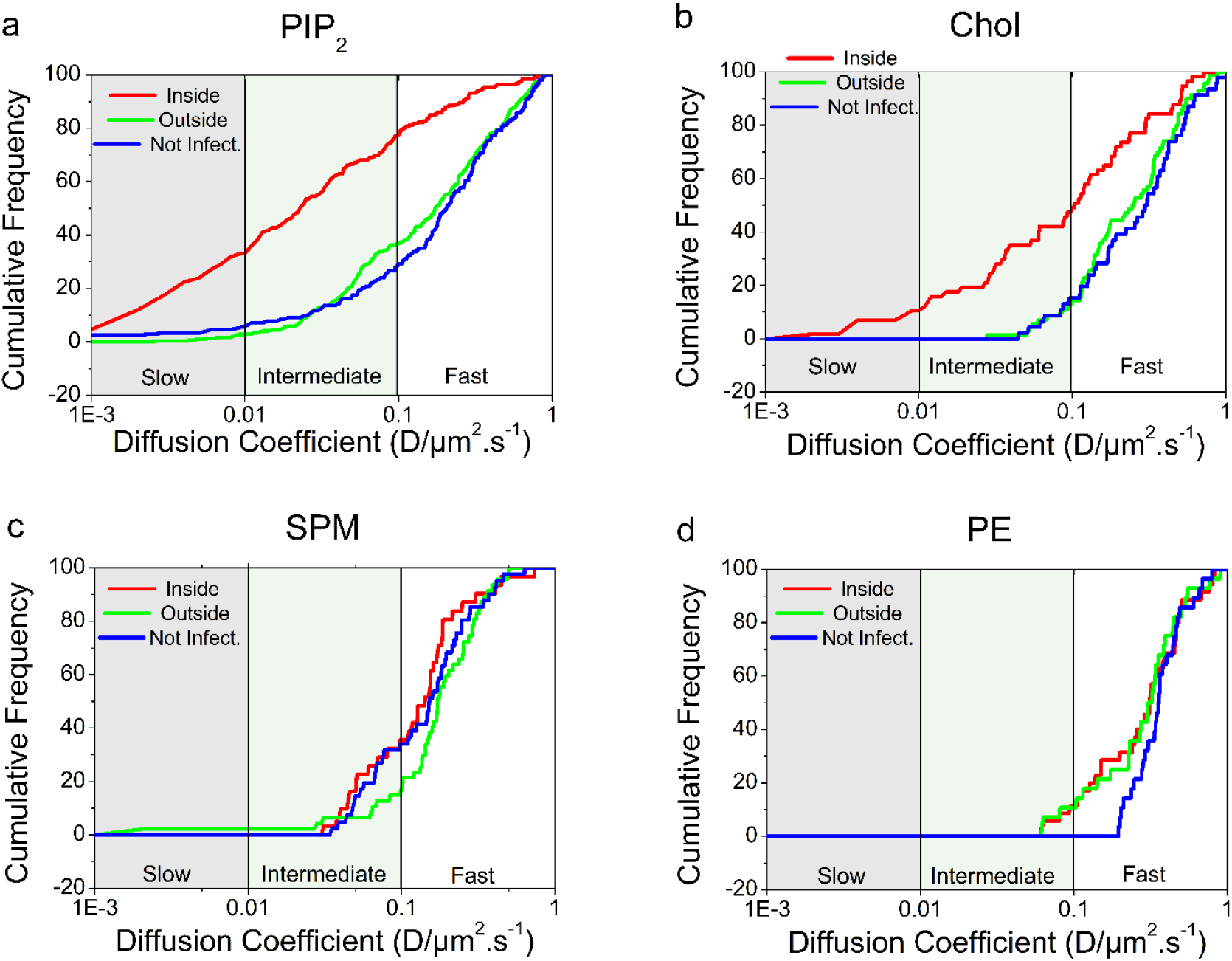
Cumulative frequency distributions of diffusion coefficient observed in HIV-1 infected T-cells. Cumulative frequency distributions of values of diffusion coefficients D measured inside (red) and outside (blue) assembly sites of infected cells, and in noninfected cells (green) for the different fluorescent lipid analogues, PI(4,5)P2 (**a**), Cholesterol (Chol) (**b**), sphingomyelin (SPM) (**c**) and PE (**d**).

### Supplementary Note 4: Cumulative frequency distributions observed in transfected cells for the different lipids

**Fig. S4.**
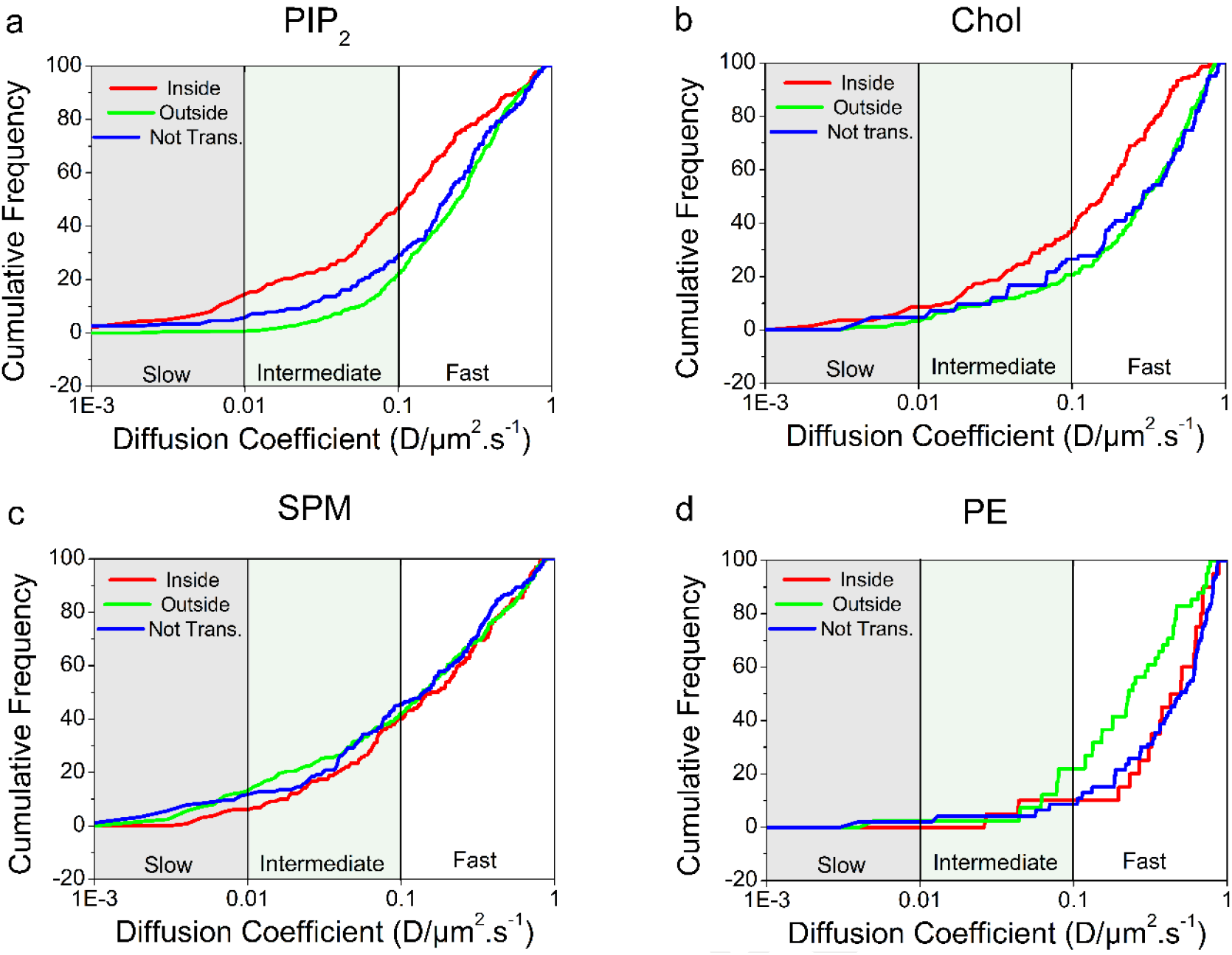
Cumulative frequency distributions of diffusion coefficient observed in HIV-1 Gag transfected T-cells. Cumulative frequency distributions of the D values measured inside (red) and outside (blue) assembly sites of transfected cells, and in nontransfected cells (green) for the different lipids, PI(4,5)P2 (**a**), Cholesterol (Chol) (**b**), sphingomyelin (SPM) (**c**) and PE (**d**).

## References

1. Britta Brügger, Bärbel Glass, Per Haberkant, Iris Leibrecht, Felix T. Wieland, and Hans-Georg Kräusslich. The HIV lipidome: a raft with an unusual composition. Proc. Natl. Acad. Sci. U.S.A., 103(8):2641–2646, February 2006. ISSN 0027-8424. doi: 10.1073/ pnas.0511136103.

2. Robin Chan, Pradeep D. Uchil, Jing Jin, Guanghou Shui, David E. Ott, Walther Mothes, and Markus R. Wenk. Retroviruses Human Immunodeficiency Virus and Murine Leukemia Virus Are Enriched in Phosphoinositides. J. Virol., 82(22):11228–11238, November 2008. ISSN 0022-538X, 1098-5514. doi: 10.1128/JVI.00981-08.

3. Jakub Chojnacki, Dominic Waithe, Pablo Carravilla, Nerea Huarte, Silvia Galiani, Jörg Enderlein, and Christian Eggeling. Envelope glycoprotein mobility on HIV-1 particles depends on the virus maturation state. Nat Commun, 8(1):545, 2017. ISSN 2041-1723. doi: 10.1038/s41467-017-00515-6.

4. Maier Lorizate, Timo Sachsenheimer, Bärbel Glass, Anja Habermann, Mathias J. Gerl, Hans-Georg Kräusslich, and Britta Brügger. Comparative lipidomics analysis of HIV-1 particles and their producer cell membrane in different cell lines. Cell. Microbiol., 15(2):292–304, February 2013. ISSN 1462-5822. doi: 10.1111/cmi.12101.

5. Robert A. Dick, Shih Lin Goh, Gerald W. Feigenson, and Volker M. Vogt. HIV-1 Gag protein can sense the cholesterol and acyl chain environment in model membranes. PNAS, 109(46):18761–18766, November 2012. ISSN 0027-8424, 1091-6490. doi: 10.1073/pnas. 1209408109.

6. Heiko Keller, Hans-Georg Kräusslich, and Petra Schwille. Multimerizable hiv gag derivative binds to the liquid-disordered phase in model membranes. Cellular microbiology, 15:237– 247, February 2013. ISSN 1462-5822. doi: 10.1111/cmi.12064.

7. Naresh Yandrapalli, Quentin Lubart, Hanumant S Tanwar, Catherine Picart, Johnson Mak, Delphine Muriaux, and Cyril Favard. Self assembly of HIV-1 Gag protein on lipid membranes generates PI(4,5)P2/Cholesterol nanoclusters. Scientific reports, 6:39332, December 2016. ISSN 2045-2322. doi: 10.1038/srep39332.

8. Balaji Olety, Sarah L. Veatch, and Akira Ono. Phosphatidylinositol-(4,5)-Bisphosphate Acyl Chains Differentiate Membrane Binding of HIV-1 Gag from That of the Phospholipase Cδ1 Pleckstrin Homology Domain. Journal of Virology, 89(15):7861–7873, August 2015. ISSN 0022-538X, 1098-5514. doi: 10.1128/JVI.00794-15.

9. Eric Barklis, August O. Staubus, Andrew Mack, Logan Harper, Robin Lid Barklis, and Ayna Alfadhli. Lipid biosensor interactions with wild type and matrix deletion HIV-1 Gag proteins. Virology, 518:264–271, May 2018. ISSN 0042-6822. doi: 10.1016/j.virol.2018.03.004.

10. Christian Eggeling, Christian Ringemann, Rebecca Medda, Günter Schwarzmann, Konrad Sandhoff, Svetlana Polyakova, Vladimir N. Belov, Birka Hein, Claas von Middendorff, Andreas Schönle, and Stefan W. Hell. Direct observation of the nanoscale dynamics of membrane lipids in a living cell. Nature, 457(7233):1159–1162, February 2009. ISSN 1476-4687. doi: 10.1038/nature07596.

11. Alf Honigmann, Veronika Mueller, Haisen Ta, Andreas Schoenle, Erdinc Sezgin, Stefan W. Hell, and Christian Eggeling. Scanning STED-FCS reveals spatiotemporal heterogeneity of lipid interaction in the plasma membrane of living cells. Nat Commun, 5:5412, November 2014. ISSN 2041-1723. doi: 10.1038/ncomms6412.

12. Aleš Benda, Yuanqing Ma, and Katharina Gaus. Self-calibrated line-scan sted-fcs to quantify lipid dynamics in model and cell membranes. Biophysical Journal, 108(3):596 – 609, 2015. ISSN 0006-3495. doi: https://doi.org/10.1016/j.bpj.2014.12.007.

13. Iztok Urbančič, Juliane Brun, Dilip Shrestha, Dominic Waithe, Christian Eggeling, and Jakub Chojnacki. Lipid composition but not curvature is a determinant of a low molecular mobility within HIV-1 lipid envelope. bioRxiv, page 315168, May 2018. doi: 10.1101/315168.

14. Balaji Olety and Akira Ono. Roles played by acidic lipids in HIV-1 Gag membrane binding. Virus Research, 193:108–115, November 2014. ISSN 0168-1702. doi: 10.1016/j.virusres. 2014.06.015.

15. Delphine Muriaux and Jean-Luc Darlix. Properties and functions of the nucleocapsid protein in virus assembly. RNA biology, 7:744–753, 2010.

16. Eric O. Freed. HIV-1 assembly, release and maturation. Nature reviews. Microbiology, 13: 484–496, August 2015. doi: 10.1038/nrmicro3490.

17. Siddhartha A. K. Datta, Xiaobing Zuo, Patrick K. Clark, Stephen J. Campbell, Yun-Xing Wang, and Alan Rein. Solution Properties of Murine Leukemia Virus Gag Protein: Differences from HIV-1 Gag. J. Virol., pages JVI.05889–11, September 2011. ISSN 0022-538X, 1098-5514. doi: 10.1128/JVI.05889-11.

18. W. Zhou, L. J. Parent, J. W. Wills, and M. D. Resh. Identification of a membrane-binding domain within the amino-terminal region of human immunodeficiency virus type 1 Gag protein which interacts with acidic phospholipids. J. Virol., 68(4):2556–2569, April 1994. ISSN 0022-538X.

19. Noémie Kempf, Viktoriia Postupalenko, Saurabh Bora, Pascal Didier, Youri Arntz, Hugues de Rocquigny, and Yves Mély. The hiv-1 nucleocapsid protein recruits negatively charged lipids to ensure its optimal binding to lipid membranes. Journal of Virology, 89(3):1756–1767, 2015. ISSN 0022-538X. doi: 10.1128/JVI.02931-14.

20. Vineela Chukkapalli, Ian B. Hogue, Vitaly Boyko, Wei-Shau Hu, and Akira Ono. Interaction between the Human Immunodeficiency Virus Type 1 Gag Matrix Domain and Phosphatidylinositol-(4,5)-Bisphosphate Is Essential for Efficient Gag Membrane Binding. J. Virol., 82(5):2405–2417, January 2008. ISSN 0022-538X, 1098-5514. doi: 10.1128/JVI.01614-07.

21. Akira Ono, Sherimay D. Ablan, Stephen J. Lockett, Kunio Nagashima, and Eric O. Freed. Phosphatidylinositol (4,5) bisphosphate regulates HIV-1 Gag targeting to the plasma membrane. Proc. Natl. Acad. Sci. U.S.A., 101(41):14889–14894, October 2004. ISSN 0027-8424. doi: 10.1073/pnas.0405596101.

22. Frauke Mücksch, Vibor Laketa, Barbara Müller, Carsten Schultz, and Hans-Georg Kräusslich. Synchronized HIV assembly by tunable PIP2 changes reveals PIP2 requirement for stable Gag anchoring. eLife Sciences, 6:e25287, June 2017. ISSN 2050-084X. doi: 10.7554/eLife.25287.

23. A. Ono and E. O. Freed. Plasma membrane rafts play a critical role in HIV-1 assembly and release. Proc. Natl. Acad. Sci. U.S.A., 98(24):13925–13930, November 2001. ISSN 0027-8424. doi: 10.1073/pnas.241320298.

24. Dzung H. Nguyen and James E. K. Hildreth. Evidence for budding of human immunodeficiency virus type 1 selectively from glycolipid-enriched membrane lipid rafts. Journal of Virology, 74(7):3264–3272, 2000. ISSN 0022-538X. doi: 10.1128/JVI.74.7.3264-3272.2000.

25. David Hawkes, Kate L. Jones, Redmond P. Smyth, Cândida F. Pereira, Robert Bittman, Anthony Jaworowski, and Johnson Mak. Properties of HIV-1 associated cholesterol in addition to raft formation are important for virus infection. Virus Res., 210:18–21, December 2015. ISSN 1872-7492. doi: 10.1016/j.virusres.2015.06.023.

26. Shahan Campbell, Katharina Gaus, Robert Bittman, Wendy Jessup, Suzanne Crowe, and Johnson Mak. The raft-promoting property of virion-associated cholesterol, but not the presence of virion-associated Brij 98 rafts, is a determinant of human immunodeficiency virus type 1 infectivity. J. Virol., 78(19):10556–10565, October 2004. ISSN 0022-538X. doi: 10.1128/JVI.78.19.10556-10565.2004.

27. Jamil S. Saad, Jaime Miller, Janet Tai, Andrew Kim, Ruba H. Ghanam, and Michael F. Summers. Structural basis for targeting HIV-1 Gag proteins to the plasma membrane for virus assembly. Proc. Natl. Acad. Sci. U.S.A., 103(30):11364–11369, July 2006. ISSN 0027-8424. doi: 10.1073/pnas.0602818103.

28. Landry Charlier, Maxime Louet, Laurent Chaloin, Patrick Fuchs, Jean Martinez, Delphine Muriaux, Cyril Favard, and Nicolas Floquet. Coarse-grained simulations of the HIV-1 matrix protein anchoring: revisiting its assembly on membrane domains. Biophysical journal, 106 (3):577–585, February 2014. ISSN 1542-0086. doi: 10.1016/j.bpj.2013.12.019.

29. Peter Y Mercredi, Nadine Bucca, Burk Loeliger, Christy R Gaines, Mansi Mehta, Pallavi Bhargava, Philip R Tedbury, Landry Charlier, Nicolas Floquet, Delphine Muriaux, Cyril Favard, Charles R Sanders, Eric O Freed, Jan Marchant, and Michael F Summers. Structural and Molecular Determinants of Membrane Binding by the HIV-1 Matrix Protein. Journal of molecular biology, 428(8):1637–1655, April 2016. ISSN 1089-8638. doi: 10.1016/j.jmb.2016.03.005.

30. Ping Chen, Wolfgang Hübner, Matthew A. Spinelli, and Benjamin K. Chen. Predominant mode of human immunodeficiency virus transfer between T cells is mediated by sustained Env-dependent neutralization-resistant virological synapses. J. Virol., 81(22):12582–12595, November 2007. ISSN 0022-538X. doi: 10.1128/JVI.00381-07.

31. Sergey Ivanchenko, William J. Godinez, Marko Lampe, Hans-Georg Kräusslich, Roland Eils, Karl Rohr, Christoph Bräuchle, Barbara Müller, and Don C. Lamb. Dynamics of HIV1 assembly and release. PLoS pathogens, 5:e1000652–e1000652, November 2009. doi: 10.1371/journal.ppat.1000652.

32. Charlotte Floderer, Jean-Baptiste Masson, Elise Boilley, Sonia Georgeault, Peggy Merida, Mohamed El Beheiry, Maxime Dahan, Philippe Roingeard, Jean-Baptiste Sibarita, Cyril Favard, and Delphine Muriaux. Single molecule localisation microscopy reveals how HIV-1 Gag proteins sense membrane virus assembly sites in living host CD4 T cells. Scientific Reports, 8(1):16283, November 2018. ISSN 2045-2322. doi: 10.1038/s41598-018-34536-y.

33. Jale Schneider, Jasmin Zahn, Marta Maglione, Stephan J Sigrist, Jonas Marquard, Jakub Chojnacki, Hans-Georg Kräusslich, Steffen J Sahl, Johann Engelhardt, and Stefan W Hell. Ultrafast, temporally stochastic STED nanoscopy of millisecond dynamics. Nature Methods, 12(9):827–830, September 2015. ISSN 1548-7091, 1548-7105. doi: 10.1038/nmeth.3481.

34. Dominic Waithe, Falk Schneider, Jakub Chojnacki, Mathias P. Clausen, Dilip Shrestha, Jorge Bernardino de la Serna, and Christian Eggeling. Optimized processing and analysis of conventional confocal microscopy generated scanning FCS data. Methods, 140-141: 62–73, May 2018. ISSN 10462023. doi: 10.1016/j.ymeth.2017.09.010.

35. A Kerviel, A Thomas, L Chaloin, C Favard, and D Muriaux. Virus assembly and plasma membrane domains: which came first? Virus research, 171(2):332–340, February 2013. ISSN 1872-7492. doi: 10.1016/j.virusres.2012.08.014.

36. Mathias J. Gerl, Julio L. Sampaio, Severino Urban, Lucie Kalvodova, Jean-Marc Verbavatz, Beth Binnington, Dirk Lindemann, Clifford A. Lingwood, Andrej Shevchenko, Cornelia Schroeder, and Kai Simons. Quantitative analysis of the lipidomes of the influenza virus envelope and MDCK cell apical membrane. J Cell Biol, 196(2):213–221, January 2012. ISSN 0021-9525, 1540-8140. doi: 10.1083/jcb.201108175.

37. Vineela Chukkapalli and Akira Ono. Molecular determinants that regulate plasma membrane association of HIV-1 Gag. J. Mol. Biol., 410(4):512–524, July 2011. ISSN 1089-8638. doi: 10.1016/j.jmb.2011.04.015.

38. Kwanyee Leung, Jae-Ouk Kim, Lakshmanan Ganesh, Juraj Kabat, Owen Schwartz, and Gary J. Nabel. HIV-1 assembly: viral glycoproteins segregate quantally to lipid rafts that associate individually with HIV-1 capsids and virions. Cell Host Microbe, 3(5):285–292, May 2008. ISSN 1934-6069. doi: 10.1016/j.chom.2008.04.004.

39. Abdul A. Waheed and Eric O. Freed. Lipids and membrane microdomains in HIV-1 replication. Virus Res., 143(2):162–176, August 2009. ISSN 1872-7492. doi: 10.1016/j.virusres. 2009.04.007.

40. Wesley I. Sundquist and Hans-Georg Kräusslich. HIV-1 assembly, budding, and maturation. Cold Spring Harb Perspect Med, 2(7):a006924, July 2012. ISSN 2157-1422. doi: 10.1101/ cshperspect.a006924.

41. Falk Schneider, Dominic Waithe, B. Christoffer Lagerholm, Dilip Shrestha, Erdinc Sezgin, Christian Eggeling, and Marco Fritzsche. Statistical Analysis of Scanning Fluorescence Correlation Spectroscopy Data Differentiates Free from Hindered Diffusion. ACS Nano, July 2018. ISSN 1936-0851. doi: 10.1021/acsnano.8b04080.

42. Jeevan B. Gc, Bernard S. Gerstman, Robert V. Stahelin, and Prem P. Chapagain. The Ebola virus protein VP40 hexamer enhances the clustering of PI(4,5)P 2 lipids in the plasma membrane. Physical Chemistry Chemical Physics, 18(41):28409–28417, 2016. ISSN 1463-9076, 1463-9084. doi: 10.1039/C6CP03776C.

43. Heidi Koldsø, David Shorthouse, Jean Hélie, and Mark S. P. Sansom. Lipid Clustering Correlates with Membrane Curvature as Revealed by Molecular Simulations of Complex Lipid Bilayers. PLoS Computational Biology, 10(10):e1003911, October 2014. ISSN 1553-7358. doi: 10.1371/journal.pcbi.1003911.

44. Marilia Barros, Frank Heinrich, Siddhartha A. K. Datta, Alan Rein, Ioannis Karageorgos, Hirsh Nanda, and Mathias Lösche. Membrane Binding of HIV-1 Matrix Protein: Dependence on Bilayer Composition and Protein Lipidation. J. Virol., 90(9):4544–4555, January 2016. ISSN 0022-538X, 1098-5514. doi: 10.1128/JVI.02820-15.

45. . Salah Edin El Meshri, Emmanuel Boutant, Assia Mouhand, Audrey Thomas, Valéry Larue, Ludovic Richert, Valérie Vivet-Boudou, Yves Mély, Carine Tisné, Delphine Muriaux, and Hugues de Rocquigny. The NC domain of HIV-1 Gag contributes to the interaction of Gag with TSG101. Biochim Biophys Acta Gen Subj, 1862(6):1421–1431, 2018. ISSN 0304-4165. doi: 10.1016/j.bbagen.2018.03.020.

46. Audrey Thomas, Charlotte Mariani-Floderer, Maria Rosa López-Huertas, Nathalie Gros, Elise Hamard-Péron, Cyril Favard, Theophile Ohlmann, José Alcamí, and Delphine Muriaux. Involvement of the Rac1-IRSp53-Wave2-Arp2/3 Signaling Pathway in HIV-1 Gag Particle Release in CD4 T Cells. Journal of virology, 89(16):8162–8181, August 2015. ISSN 1098-5514. doi: 10.1128/JVI.00469-15.

47. Mathias P. Clausen, Erdinc Sezgin, Jorge Bernardino de la Serna, Dominic Waithe, B. Christoffer Lagerholm, and Christian Eggeling. A straightforward approach for gated sted-fcs to investigate lipid membrane dynamics. Methods, 88:67 – 75, 2015. ISSN 1046-2023. doi: https://doi.org/10.1016/j.ymeth.2015.06.017. Super-resolution Light Microscopy.

